# Heterochromatin Replication: Direct Interaction of DNA replication machinery with heterochromatin code writer Clr4/Suv39 and reader Swi6/HP1 in *S. pombe*

**DOI:** 10.1101/2020.10.21.349183

**Authors:** Sharanjot Saini, Sumit Arora, Kamlesh K. Bisht, Nandni Nakwal, Shakil Ahmed, Jagmohan Singh

## Abstract

The establishment of heterochromatin in fission yeast involves methyltransferase Clr4-mediated H3-Lys9 methylation, which is bound specifically by Swi6/HP1. However, the mechanism of propagation of heterochromatin through multiple cell divisions is not known. A role of DNA replication in propagating the heterochromatin is envisaged. Studies in *S. pombe* have indicated a direct interaction between DNA Polα and Swi6/HP1 and between DNA Polε and Rik1-Dos2 complex, suggesting a coupling between DNA replication and heterochromatin assembly. Here, we show that like DNA Polα, Polδ, which plays a role in both leading and lagging strand replication, also plays a role in silencing at mating type and centromere. We show that both the polymerases α and δ interact directly with both Clr4 and Swi6/HP1. Mutations in both the polymerases lead to decrease in H3-Lys9 methylation and Swi6 at the mating type and left outer repeats of centromeres I and II, with a reciprocal increase in their level at the central element, *cnt*, at all the three centromeres. These mutations also cause defects in chromosome segregation, recruitment of Cohesin and chromosome dynamics during mitosis and meiosis. Thus, our results indicate that a tight coordination between DNA replication machinery and propagation of the heterochromatin-specific epigenetic mark.

---

Propagation of state/s of expression of genes through multiple cell divisions underlies the process of differentiation and development in all eukaryotes, including the unicellular eukaryotes, like the budding and fission yeasts, Drosophila, mouse and humans. Notable insights in recent years have revealed that more than DNA, it is the chromatin structure which functions as a Mendelian unit of inheritance (1). In an elegant study, Grewal and Klar (2) showed that in fission yeast two alternative epigenetic states of the mating type locus were propagated through not only multiple mitotic divisions but segregated as alternative Mendelian alleles during meiosis. Similar findings of stable epigenetic chromosomal states have been reported in other systems (3). Subsequent work in fission yeast has revealed that while RNAi machinery is required for the establishment of silencing, the heterochromatin proteins Swi6/HP1 and the methyltransferase Clr4 are involved in spreading of the heterochromatin (4).

However, the mechanism of propagation of epigenetic states during multiple cell divisions is not well understood. Earlier studies in *S. cerevisiae* showed a requirement of passage through S phase for establishment of mating-type silencing (5), which were supported by later findings showing the requirement of a functional replication origin and the proteins, that assemble the Origin Recognition Complex at replication origins, in silencing (7–10). However, later, silencing was shown to occur on an artificial template without undergoing replication (11,12). Further, a role of Cohesin degradation during mitosis in silencing suggests that the existence of more than one control in establishment and propagation of silencing in *S. cerevisiae* (13,14).

Despite these observations, the idea that DNA replication may play a pivotal role in propagation of heterochromatin structure remains intuitively attractive because of the implicit simplicity and ability to explain the faithful assembly of heterochromatin through direct interaction between DNA replication and heterochromatin machineries. In fact, work from our laboratory and others has shown a role of DNA polymerase α in propagation of heterochromatin at the mating type, centromere and telomere loci through recruitment of the chromodomain protein Swi6/HP1 in *S. pombe* (15–17). Accordingly, we proposed a replication-mediated recruitment model for propagation of heterochromatin structure (15), where in the replication enzyme, DNA Polα, which is needed for carrying out lagging and leading strand synthesis in all eukaryotes, may recruit the chromodomain protein Swi6/HP1 coincidentally with DNA replication (15–17). In another study, interaction of Polε, which is also involved in DNA replication, with the Rik1-Dos2 complex, was shown to play a role in heterochromatin silencing (18). Here, we demonstrate that both DNA Polα and Polδ, in addition to their role in coordinating the lagging and leading strand replication in eukaryotes, also bind to and help in recruiting the methyltransferase Clr4 and chromodomain protein Swi6/HP1 (which form the dual components of histone code involved in writing the heterochromatin specific code of H3-Lys9 methylation and reading of this code, respectively; 19) to the mating type and centromere loci. Both *polα* and *polδ* mutants are defective in silencing at the mating type and centromere loci; this effect is related to reduced localization of Swi6/HP1 and H3-Lys9 methylation at these loci. The mutants also exhibit defective chromosome cohesion due to lack of Cohesin binding, an effect related to defective recruitment of Swi6/HP1, as well as other defects like sensitivity to cold and microtubule destabilizing drug thiabendazole and aberrant chromosome segregation during mitosis and meiosis. Surprisingly, the mutants also show enhanced localization of Swi6 and H3-Lys9-Me2 at the central region, *cnt*, of all three centromeres in place of the centromeric histone, Cnp1/CENPA. Thus, the components of leading and lagging strand replication orchestrate fidelity of replication of heterochromatin regions through direct interaction with and recruitment of the heterochromatin proteins, thus integrating DNA replication with the processes of silencing, sister chromatid cohesion and chromosomal dynamics during mitosis and meiosis.

## MATERIALS AND METHODS

### Strains and media

The list of strains is given in Table 1 (Supplementary section). Media and conditions were as described earlier (20). Most strains were grown at 25°C. The strains used for silencing assays carried a non-switchable *mat1* locus, *mat1Msmto* or *mat1P 17::LEU2* and the donor loci *mat2* and *mat3*, whose cis -acting repression elements *REII* and *REIII* were deleted along with insertion of *ura4* and *ade6* reporters, respectively, as shown in Figure 1A. The two-hybrid system was a gift of Stephen Elledge (21).

**Figure 1.**
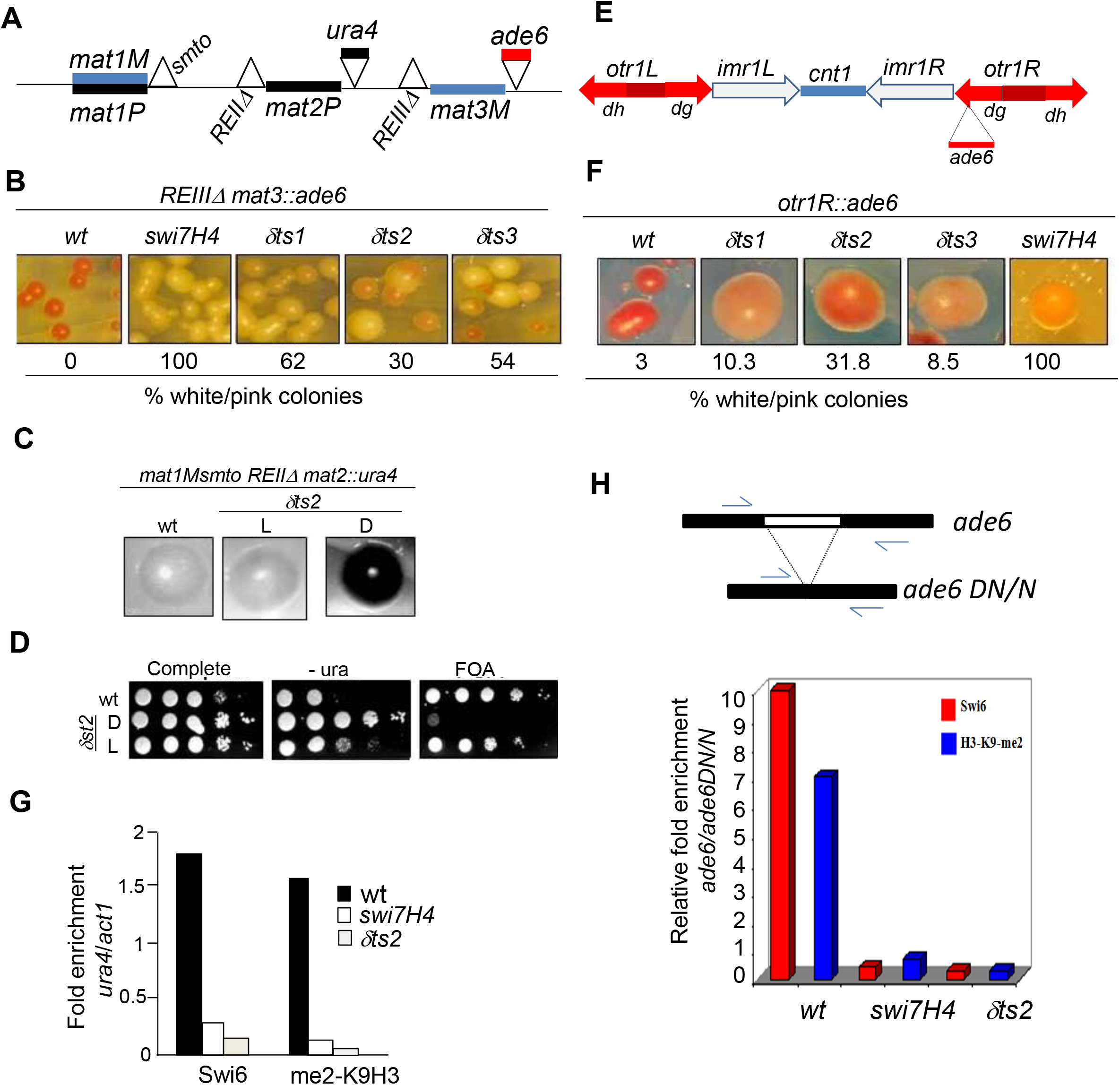
*polα* and *polδ* mutations abrogate silencing and localization of Swi6 and H3-Lys9 methylation at mating type and centromere loci. (A) The organization of mating type locus, depicting the active *mat1* locus and the silent loci *mat2* and *mat3*. Sites of deletion of the cis-acting silencers *REII* and *REIII*, flanking *mat2* and *mat3* loci, respectively, are indicated by the sign. Also indicated are the insertions of *ura4* near *mat2* and *ade6* reporter gene near the *mat3* locus. (B) Both *polα (swi7H4)* and *polδ* mutations (*δts1-δts3*) cause derepression of the *ade6* reporter linked to *mat3* locus. The colonies of the wild type and mutant strains were streaked on adenine limiting plates YE (18), and grown at 25°C for 4 days (top panel). (C) *δts2* mutation causes derepression of the silent locus *mat2* having its repression element deleted (*REII*). Cells of the indicated strains were streaked on PMA plates, grown for 3-4 days at 25°C and stained with iodine. Cells of dark and light staining colonies were cultured, serial dilutions spotted on complete, -ura and FOA plates and allowed to grow at 25°C for 3-4 days. (D) ChIP assay was performed with the strains having the *REII* deleted in wild type, *swi7H4* and *δts2* mutant backgrounds and enrichment of Swi6 and Lys9-methylated H3 at *ura4* reporter linked to *mat2* locus versus *act1* as a negative control quantitated. Histogram shows the quantitation of the ChIP data representing the average of two independent experiments. (E) A schematic representation of the *cenI* locus, showing insertions of *ura4* reporter at the *cnt1* and *imr1L* loci and insertion of *ade6* reporter at the outer repeat *otr1R*. (F) *polδ* mutations derepress the *ade6* reporter at the outer repeat *otr1R* of *cenI*. The indicated strains were streaked on adenine-limiting plates (YE), allowed to grow at 25°C and photographed. The percentage of pink and white colonies are indicated. (G,H) *polα* (*swi7H4*) and *polδ* (*δts2*) mutations cause depletion of Swi6 and H3-Lys9 methylation at the *mat2::ura4* (G) or *ade 6* reporter inserted at the *otr1R* region of *chrI* (H). Strains of the indicated strains were subjected to ChIP assay and enrichment of Swi6 and H3-Lys9 methylation at the *ura4* for *mat2*-linked *ura4* reporter versus *act1* (G) and *ade6* located at the heterochromatic *otr1R* versus the euchromatic *ade6DN/N* (H) quantitated by PCR. Histogram shows the quantitation of the ChIP data representing the average of two independent experiments.

### Plasmsid constructs

The construct expressing MBP-tagged Polα was described earlier (15). The GST-tagged Polδ construct, pGST-pol3-N 1-211, expressing truncated *polδ* gene lacking the first 211 codons cloned into pGEX4T-1, was a gift from Dr. S. MacNeill (22). To construct the plasmid expressing (His)_6_-tagged, *clr4* gene was PCR amplified using the primers Clr4-For Bam (ATGCGGATCCTCGCCTAAACAAGAGGAGTAT) and Clr4 RevHind (ATGCAAGCTTTTAACCGAAAAGCCAG CCAC). The PCR product was cleaved with *Bam*HI and *HindIII* and cloned into the vector pQE30 which was cleaved with the same enzymes. The construct to express (His)_6_-tagged Swi6 was prepared by amplifying the *swi6* gene using the primers Swi6-N-Bam (ATGCGGATCCCAAGAAAGGAGGTGTTCG) and Swi6-C-Rev-Hind (ATGCAAGCTTATTTTCACGGAACGTTA AG). The PCR product was cleaved with BamHI and HindIII and cloned into the vector pQE30, which was cleaved with the same enzymes.

### Silencing assays

Expression of *ura4* reporter inserted at various heterochromatin loci in wild type and mutant strains was monitored by the serial dilution spotting assays. 5μl aliquots of 10-fold serial dilutions of cultures of the strains were spotted on complete plates or plates lacking uracil or containing FOA. Plates were incubated at 25°C for 4-5 days. Expression of the *ade6* reporter inserted at the *otr1R* region of *cenI* was monitored by streaking the cells on YE plates and incubating at 25°C for 4-5 days. While the wild type strains harbouring the *ade6* locus in the repressed state at the *mat* or centromere loci appear red, the mutants yield white or pink colonies. For monitoring silencing in the silencer deletion background, strains were also streaked on PMA^+^ plates (20). After growth at 25°C for 5 days they were either stained with iodine or their cells were examined microscopically for the haploid meiosis phenotype, which results from simultaneous expression of the Minus and Plus transcripts in the silencing mutant (20).

### Pull-down assays

The GST-Clr4 fusion protein was allowed to bind to Glutathione-agarose beads (Sigma) in MTPBS buffer (100mM Na_2_HPO_4_, 16mM NaH_2_PO_4_, 150mM NaCl). Glutathione-agarose beads were equilibrated in MTPBS buffer before use. The fusion protein was added to the reaction mix containing 1mg/ml BSA, 50μl of beads (50% v/v) in MTPBS buffer and allowed to bind for 1hr at 4°C with gentle mixing. Reaction with GST was included as a control and the reaction with the fusion protein was set up in triplicate. After binding, the GST-Clr4 fusion protein bound beads were collected by centrifugation (500g for 10 sec) and washed twice with MTPBS. Following the washings, three increasing concentrations of MBP-Polα were added in addition to 1mg/ml BSA and 1X binding buffer [5X binding buffer 750mM NaCl, 100mM Tris-Cl (pH 8.0), 5mM EDTA, 0.5% Triton X-100, 5mM DTT]. The reactions were incubated by gentle mixing for 2 hours at 4°C followed by four washings with the wash buffer (100mM NaCl, 20mM Tris-Cl (pH 8.0), 1mM EDTA, 0.1% Triton X-100, 1mM DTT). After washings, the beads were suspended in 1X SDS loading dye. Samples were boiled for 10 minutes and resolved by SDS-PAGE along with the input material.

In other pull-down experiments, extracts were prepared from *E. coli* strains expressing (His)_6_-Clr4 and (His)_6_-Swi6 and allowed to bind at 4°C overnight to Ni-NTA resin (Qiagen; binding capacity 5-10mg protein/ml resin), which was equilibrated with binding buffer [50mM NaH_2_PO_4_ (pH 8.0), 300mM NaCl, 10mM imidazole]. Washings were performed 3 −4 times with the same buffer at 4°C and the beads were collected by centrifugation at 500g for 5 min. Now the crude extract prepared from strains expressing GST-Polδ or GST-Polα was added to the above resin at increasing concentrations (from 100 to 300μg) along with control, 1mg/ml BSA and 1X binding buffer (5X binding buffer: 750 mM NaCl, 100mM Tris-Cl, pH 8.0, 5mM EDTA, 5mM DTT) was added, incubated at 4°C for 3-4 hours to overnight and then washed four times with washing buffer (100mM NaCl, 20mM Tris-Cl, 1mM EDTA, 1mM DTT). To the above samples 1X SDS loading buffer was added and samples boiled for 5-10 min. The samples were subjected to SDS-PAGE, Western blotted and probed with anti GST antibody.

### Chromatin Immunoprecipitation (ChIP) assay

ChIP assay was performed as described earlier (23). The following oligos were used for PCR: ura4F: gaggggatgaaaaatcccat; ura4R: ttcgacaacaggattacgacc; ade6F: tgcgatgcacctgaccaggaaagt; ade6R: agagttgggtgttgatttcgctga;act1F: tcctacgttggtgatgaagc; act1R: tccgatagtgataacttgac; dhFor: ggagttgcgcaaacgaagtt; dhRev: ctcactcaagtccaatcgca. PCR conditions were: 95°C, 2 min; 30 cycles of 95°C, 1 min-55°C, sec-72°C 2 min; 72°C, 10 min. 0.1 μCi of (α^32^-P)-dCTP was included during PCR and the PCR products were resolved by polyacrylamide gel electrophoresis, followed by quantitation with BioRad phosphoimager.

### Confocal microscopy

Sporulating asci and vegetative cells were stained with the nuclear stain 4’, 6’-diamidino-2-phenylindole (DAPI) to visualise the meiotic and/or mitotic division defects in *swi7H4* and *δts* mutants. The 10X DAPI stock was diluted with glycerol to 4X just prior to use. For studying the meiotic defects, cells were plated on PMA^+^ plates and were grown for 3-4 days at 25 °C to allow an adequate level of sporulation. Sporulating asci were scraped off the plate and suspended in 1X PBS. The cells or spores were fixed by mixed aldehyde method (Paul Nurse’s lab protocols: ‘Fission Yeast Handbook’), which involves the addition of fresh 37% paraformaldehyde solution to a final concentration of 3.7%, followed by the addition of glutaraldehyde to 0.2% final concentration. Fixation was allowed to proceed for 60-90 min at RT with gentle shaking. After fixation, cells and asci were collected by centrifugation, washed thrice with 1X PBS, followed by resuspension in PBS-T (PBS + Triton X-100). After 30 sec in PBS-T, cells were washed thrice in 1X PBS. After the final wash, cells were resuspended in 100-150μl 1X PBS. 6μl of this cell suspension was mixed with 4μl of 4X DAPI on a microscopic slide and coverslip was gently placed on the slide, pressing with a tissue paper to remove excess liquid. The samples were then visualized with a Carl Zeiss LSM 510 Meta confocal microscope.

For studying the chromosome dynamics during mitosis, vegetative cells were cultured in 5ml YEA (20) overnight at 25°C. Cells were then reinoculated into 10-15ml fresh media (OD_600_ 0.1-0.2) and incubated at 18°C with shaking till mid-log phase (OD_600_ 0.5-0.6). Cultures were then harvested (6000 rpm for 2 min). Cells were then fixed with methanol by initially resuspending in 1ml methanol (−20°C) followed by the addition of ~9ml chilled methanol. Cells were then kept at −20°C for 10 minutes to overnight. After fixation, cells were stained with DAPI as well as with α-tubulin antibody, TAT1 to visualize the mitotic spindle along with the nuclei.

### ChlP-ChIP Analysis

ChIP-samples used to measure the distribution of H3-K9-Me2 and Swi6 at centromeric regions according to Ekwall and Partridge (23), were processed for ChIP-on-ChIP analysis.according to Sandmann *et al.* (24), using Affymetrix non-redundant microarrays.

## RESULTS

### Mutations in Polα and Polδ abrogate silencing at the mating type and centromere loci

To extend our earlier work showing a role of DNA Polα in silencing, we checked whether DNA replication machinery, in general, is required for silencing, as it has been shown that both Polα and Polδ play an important role in both leading and lagging strand replication (25, 26). Accordingly, we tested the effect of mutations in *polδ*, namely *δts1, δts2* and *δts3* (27) on silencing at the mating type locus. The genetic screen employed a strain in which the repression elements flanking the silent *mat* loci, *REII* flanking the *mat2* locus and *REIII* flanking the *mat3* locus (Figure 1A) were deleted along with insertion of *ura4* and *ade6* reporters, respectively. Earlier studies showed that mutations in *swi6* and *clr1-clr4* abrogated silencing strongly when combined with deletion of the repression elements (28,29). Mutations in *polα (swi7H4)* and *polδ* increased expression of the *ade6* reporter flanking the *mat3* locus, as indicated by growth of light pink colonies instead of the red colonies of the parent strain, which is indicative of loss of repression in the mutant (Figure 1B). Further, as shown earlier in case of the *polα* mutant *swi7H4* (15), *δts2* mutant showed similar loss of silencing of the *mat2P* locus, as indicated by iodine staining of the colonies (15; see Materials and Methods; Figure 1C). Interestingly, like *swi7H4/polα* mutant (16), *polδts2* mutant also exhibited two epigenetic states that give light (L) and dark (D) staining with iodine, representing the repressed and derepressed state of the *mat2P* locus, respectively (Figure 1C). These states also show, respectively, low and high level of the expression of the *mat2P*-linked *ura4* reporter, as indicated by level of growth on plates lacking uracil (Figure 1D). These results indicate that like Polα (15–17), Polδ is also involved in the same pathway of silencing as Swi6 and Clr1-Clr4 and plays a role in the establishment of the repressed state.

We showed earlier that *swi7H4/polα* mutation also abrogated silencing at the centromere loci. Therefore, we tested the effect of *δts1-δts3* mutations on silencing at the centromere loci by monitoring the expression of the *ura4* reporter inserted at the *cnt1* and *imr1* regions and the *ade6* reporter at the outer repeat *otr1R* (Figure 1E). Results showed that the mutations did not affect silencing at the *cnt1* and *imr1L* region (not shown). On the other hand, silencing of the *ade6* reporter inserted at the *otr1R* repeat was abrogated, as indicated by growth of pink and white colonies (10-34% of total colonies) in all the *polδ* mutants as compared to red colonies in wt strain on adenine limiting media (Figure 1F). However, unlike the *polα* (15,17) there was no effect of *polδ* mutations on silencing at the telomere loci (not shown).

### Reduced heterochromatin localization of Swi6/HP1 and H3-Lys9 methylation in *polα* and *polδ* mutants

As silencing is associated with localization of Swi6 and H3-Lys9-Me2 to heterochromatin loci, we checked the localization of Swi6 and H3-Lys9-Me2 to the *mat* and *cen* loci by Chromatin immunoprecipitation (ChIP) assay. Results showed that localization of Swi6 as well as H3-Lys9 methylation was drastically reduced at the *ura4* reporter flanking the *mat2* and *ade6* reporter flanking *otr1R* repeat of *cenI* in both *swi7H4/polα* and *δts2* mutants (Figure 1G, 1H). These results were paralleled by those of ChIP-on-ChIP experiment, wherein, especially the left *otr1* regions of *chrI* and *chrII* showed a similar reduction in the level of H3-Lys9-Me2 and Swi6 (Figure 2, double vertical arrow heads; Supplementary Figures 1 and 2)

**Figure 2.**
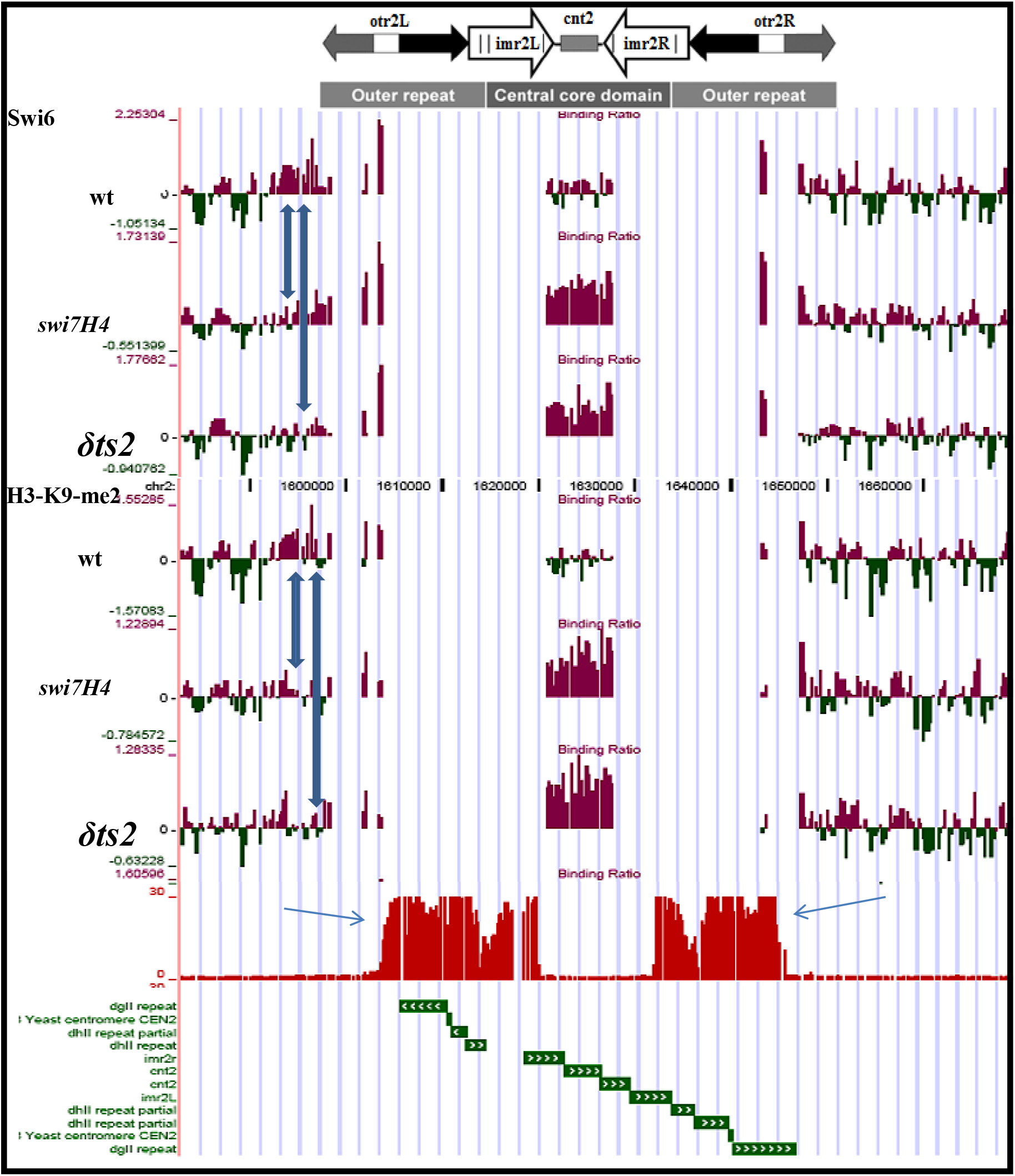
ChIP-on-ChIP assay to quantitate the level of H3-Lys9-Me2 and Swi6 in wild type versus *swi7H4/polα* and *polδts2* mutants at the *cenI*. Double vertical arrows indicate the regions in the outer repeat reduction in the level of H3-Lys9-Me2 and Swi6.

Surprisingly, results of ChIP-on-ChIP also revealed an enhanced localization of both Swi6 and H3-Lys9-me2 at the *cnt* region of Chromosome I (Figure 2) as well as *chr*II (Supplementary Figure 1) and *chr*III (Supplementary Figure 2), which is normally occupied by the centromere-specific histone H3 variant, *cnp1*/CENPA..

Earlier work showed that GFP-tagged Swi6 is localized at mainly three foci, representing the heterochromatin regions, in majority of cells (30). In fact, *swi7H4/polα* as well as *swi7-1* mutations were shown to cause delocalization of GFP-tagged Swi6 (15,17). Therefore, we determined the localization of Swi6 in *polδ* mutants by confocal microscopy. Results showed that, as shown in *swi7H4/polα* mutant earlier (15), the distribution of GFP-Swi6 was drastically reduced in *polδ* mutants, with a larger percentage of cells having only one or two foci. (Supplementary Figure 3A).

Silencing proteins like Swi6 have also been shown to be required for microtubule stability. Thus, *swi6* mutant exhibits enhanced sensitivity to cold (18-20°C) and the microtubule destabilizing drug, thiabendazole (30,31). We tested the same properties in case of *swi7H4/polα* and *δts1-δts3* mutants. Interestingly, all the mutants show a level of sensitivity to cold and thiabendazole similar to that observed with *swi6* mutant (Supplementary Figure 3B). Thus, both the polymerases also promote the mitotic microtubule stability and fidelity of chromosome segregation, which may be ascribed to their role in recruitment of Swi6 and H3-Lys9-Me2.

### Direct physical interaction of DNA Polα and Polδ with Swi6 and Clr4

The above results suggested the possibility that Polα and Polδ may interact with Swi6 and/or Clr4. Earlier work had shown that Polα binds to Swi6 *in vitro* and *in vivo* (15,17). We carried out pull-down assay to determine the binding of GST-tagged Clr4 to MBP-tagged Polα. Results showed that GST-Clr4 was retained by amylose beads to which MBP-Polα was bound (Figure 3A, lanes 3-5) but not to beads on which MBP was immobilized (Figure 3A, lane 7). It was shown earlier that mutant Polα does not interact with Swi6 *in vitro* (15, 17). Pull-down experiment using extracts from wt and *swi7H4* mutant cells showed that interaction of mutant Polα with Clr4 was at least 3-fold lower than that observed with wt Polα (Figure 3B). Furthermore, results of 2-hybrid experiment, supported the possibility of interaction of Polα with Clr4 *in vivo* (Figure 3C, 3D)

**Figure 3.**
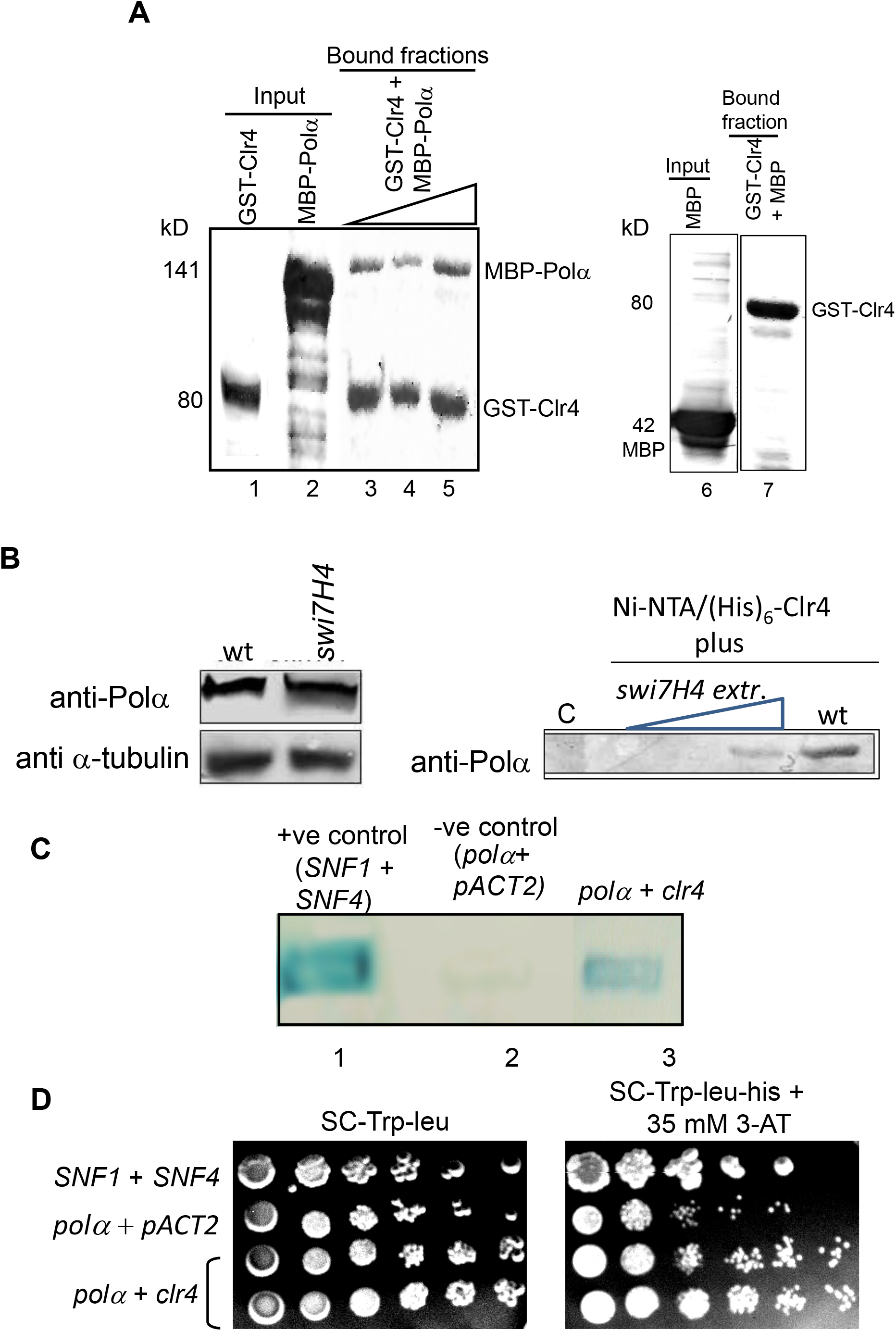
Interaction of Clr4 with DNA Polα *in vitro and* in vivo. (A) Polα interacts with Clr4 in *vitro*. Pull-down assay was performed in which GST-Clr4 was immobilized on Gluthione agarose beads and increasing concentrations of MBP-Polα (lanes 3-5) were bound to the beads. The bound fractions were resolved by SDS-PAGE and stained with Coomassie blue. In another control MBP was reacted with the beads to which GST-Clr4 was bound (lane 7), showing no MBP indicating lack of binding of GST-Clr4. (B) Reduced binding of mutant *Polα/swi7H4* protein to Clr4 *in vitro*. Increasing concentrations of extracts of *swi7H4* mutant cells were treated with Ni-NTA beads to which extract prepared from the *E. coli* strain expressing (His)_6_-tagged Clr4 was bound. C represents the Ni-NTA beads alone to which wt extract was bound. Wild type extract was added at the same concentration as the highest concentration of *swi7H4* mutant extract 300μg). (C, D) Two-hybrid assay showing interaction between Polα and Clr4 *in vivo*. Shown is the ability of two hybrid system co-expressing *polα* and *clr4* to activate the expression of lacz (C) or *his3* reporter gene (D).

We further checked whether Swi6 and or Clr4 bind to Polδ as well. Results of pull-down assay revealed that GST-tagged Polδ (Figure 4A, lanes 3-5), but not GST (Figure 4A, lane 8), could be specifically retained on the Ni-NTA beads on which (His)_6_-tagged Swi6 (Figure 4A, lanes 3-5) as well as (His)_6_-tagged Clr4 was immobilized (Figure 4B, lanes 3-5; lane 8). Furthermore, antibodies against myc epitope and Swi6 could coimmunoprecipitate Swi6 along with myc-tagged Polδ in wt but not in *swi6Δ* strain (Figure 4C). Thus, Polα and Polδ bind to both Swi6 and Clr4 *in vitro* and *in vivo* and the inability of *polα* (and possibly *polδ*) mutant proteins to recruit Swi6 and H3-Lys9 methylation may be correlated with their reduced binding to Swi6 and Clr4.

**Figure 4.**
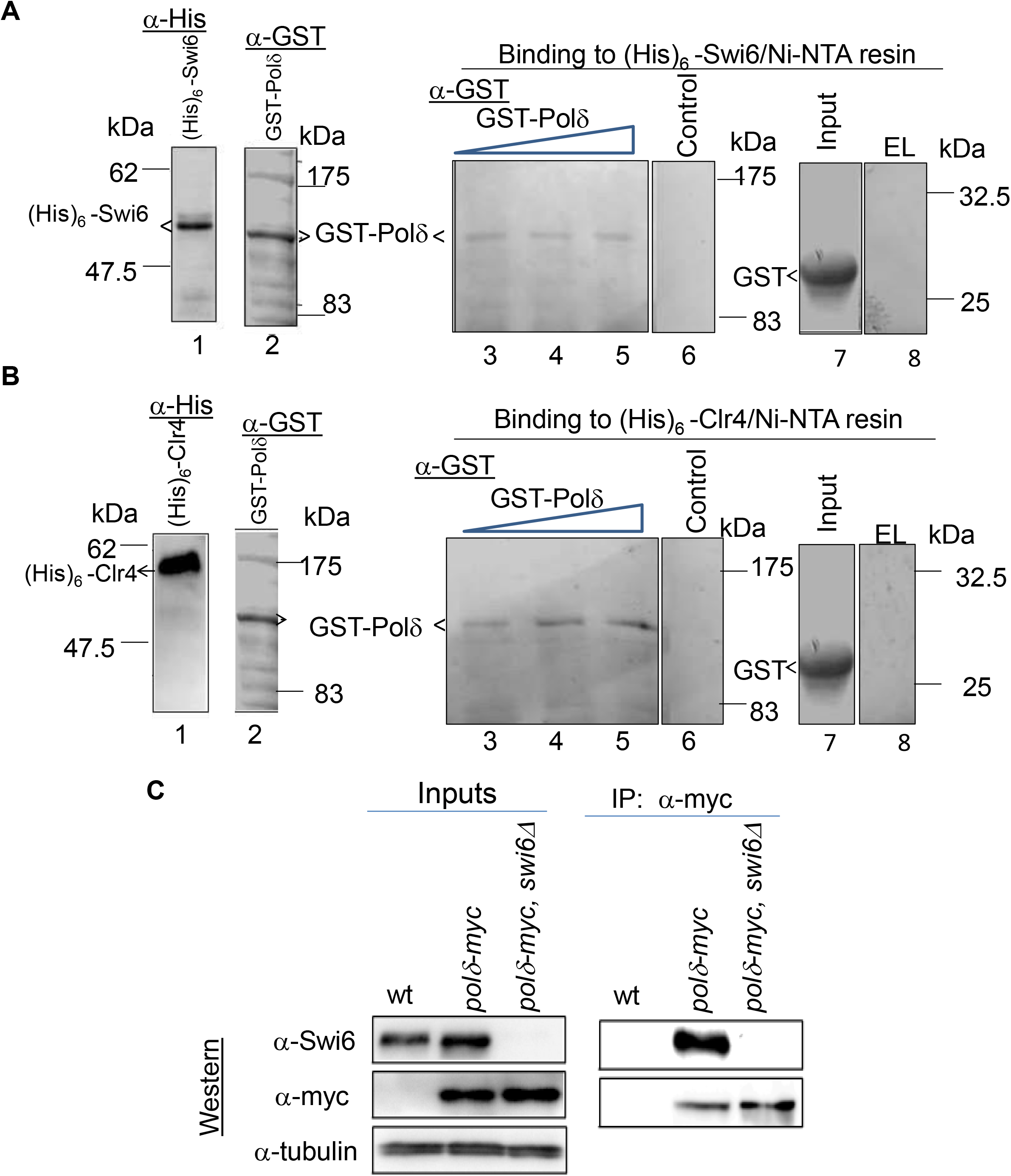
Polδ interacts with Swi6 and Clr4 *in vitro* and *in vivo*. (A) Pull-down assay was performed by binding increasing concentrations of GST-Polδ to Ni-NTA resin on which (His)_6_-tagged Swi6 was immobilized (lanes 3-5). The control represents Ni-NTA column alone (lane 6). Lanes 1 and 2 represent the loading controls of (His)_6_-Swi6 and GST-Polδ, respectively. As controls, binding of GST alone (lane 7) to Ni-NTA beads was tested (lane 8). (B) Pull-down assay was performed by binding increasing concentrations of GST-Polδ to Ni-NTA resin on which (His)_6_-tagged Clr4 was immobilized (lanes 3-5). The control represents Ni-NTAcolumn alone (lane 6). Lanes 1 and 2 represent the loading controls of (His)_6_-Clr4 and GST-Polδ, respectively. As controls, binding of GST alone (lane 7) to Ni-NTA beads was tested (lane 8). The bound fractions were resolved by SDS-PAGE and immunoblotted with anti-GST antibody. (C) Interaction of Polδ with Swi6 *in vivo*. Extracts from untagged and *polδ-myc* tagged in *swi6^+^* or *swi6Δ* background were immunoprepitated followed by western blotting with anti-myc and Swi6 atnibodies.

### Abrogation of Cohesin binding, sister chromatid cohesion and microtubule stability in *polα* and *polδ* mutants

In addition to the effect on silencing, Swi6/HP1 has also been shown to be necessary for recruitment of Cohesin to the centromere and mating type loci (32,33) and thus instrumental in enhancing the stability of chromosomes and facilitating faithful chromosome segregation. Since *swi6* mutant has been shown to be defective in sister chromatid cohesion due to lack of recruitment of Cohesin, we tested the phenotypes of *swi7H4/polα* and *δts1-δts3* mutants, both of which are defective in Swi6 recruitment. Sister chromatid cohesion can be monitored microscopically with the help of a strain system harboring LacI-GFP fusion protein and containing the LacO DNA repeat inserted at the *lys1* locus linked to centromere 1 (*cen*1-GFP) (34). The binding of the GFP-tagged repressor to the repeats appears in the form of one single dot in G2 cells representing the two sister chromatids being held tightly together at the centromere region (34). Thus, nearly all wild type cells show one spot of GFP, while *swi7H4, δts* mutants show a high fraction of cells (9-27%) having two GFP spots, indicating a loss of sister chromatid cohesion in the mutants (Supplementary Figure 4A).

To directly check whether the binding of cohesin to the centromere loci was affected, we carried out ChIP assay. Results of ChIP analysis with strains carrying HA-tagged copy of the Rad21 subunit of cohesin showed that localization of Rad21, which is highly enriched in the *dh* repeats in the *otr1R* region (32,33), is severely abrogated in *swi7H4/polα* and *δts2* mutants (Supplementary Figure 4B).

### Enhanced chromosomal segregation defects during mitosis in *polα* and *polδ* mutants

Proper assembly of heterochromatin has been shown to be important not only for silencing but also for chromosome segregation. Microscopic examination of mitotic nuclei has shown that while in wild type the two daughter nuclei of equal sizes segregate in an equidistant fashion among the presumptive daughter cell halves during cell division, the mutants defective in silencing, like swi6, *clr4* and *rik1*, show aberrant segregation of daughter nuclei. Pidoux *et al.* (35) had classified the various types of lagging chromosome segregation defects. As we analyzed only fixed cells not live cells as in case of Pidoux *et al.* (35), for convenience, we have redefined the classes of defects based on but slightly differently from ref 33. Following detailed observation of dividing cells of *swi7H4/polα* and *δts1-δts3* mutants, we identified various types of segregation defects (Figure 5A and 5B). Results show that all the *polδ* mutants exhibit the lagging chromosome phenotype (class I): while *δts1, δts3* and *swi7H4* show the lagging chromosome phenotype with a chromatid being unable to catch up with the daughter nucleus (class II; Figure 5A), class II defect of lagging chromosomes in the context of uneven segregation was observed in *δts3* and *swi7H4* mutants. A grossly defective segregation resulting in fragmented nuclei (class V) was observed in *δts2* and *δts3* mutants, while unequal movement of daughter nuclei towards the poles (class VI) was observed only in *δts2* mutant (Figure 5A). The total percentage of cells showing aberrant segregation defects is depicted in Figure 5B. Thus, both *polα* and *polδ* mutants exhibit gross chromosomal segregation defects, that are reminiscent of those exhibited by *swi6* mutant (33). It is noteworthy that microtubule integrity is also severely compromised in the mutants showing discontinuous, fragmented and punctate pattern, unlike a straight, uninterrupted pattern observed in wild type cells (Figure 5A).

**Figure 5.**
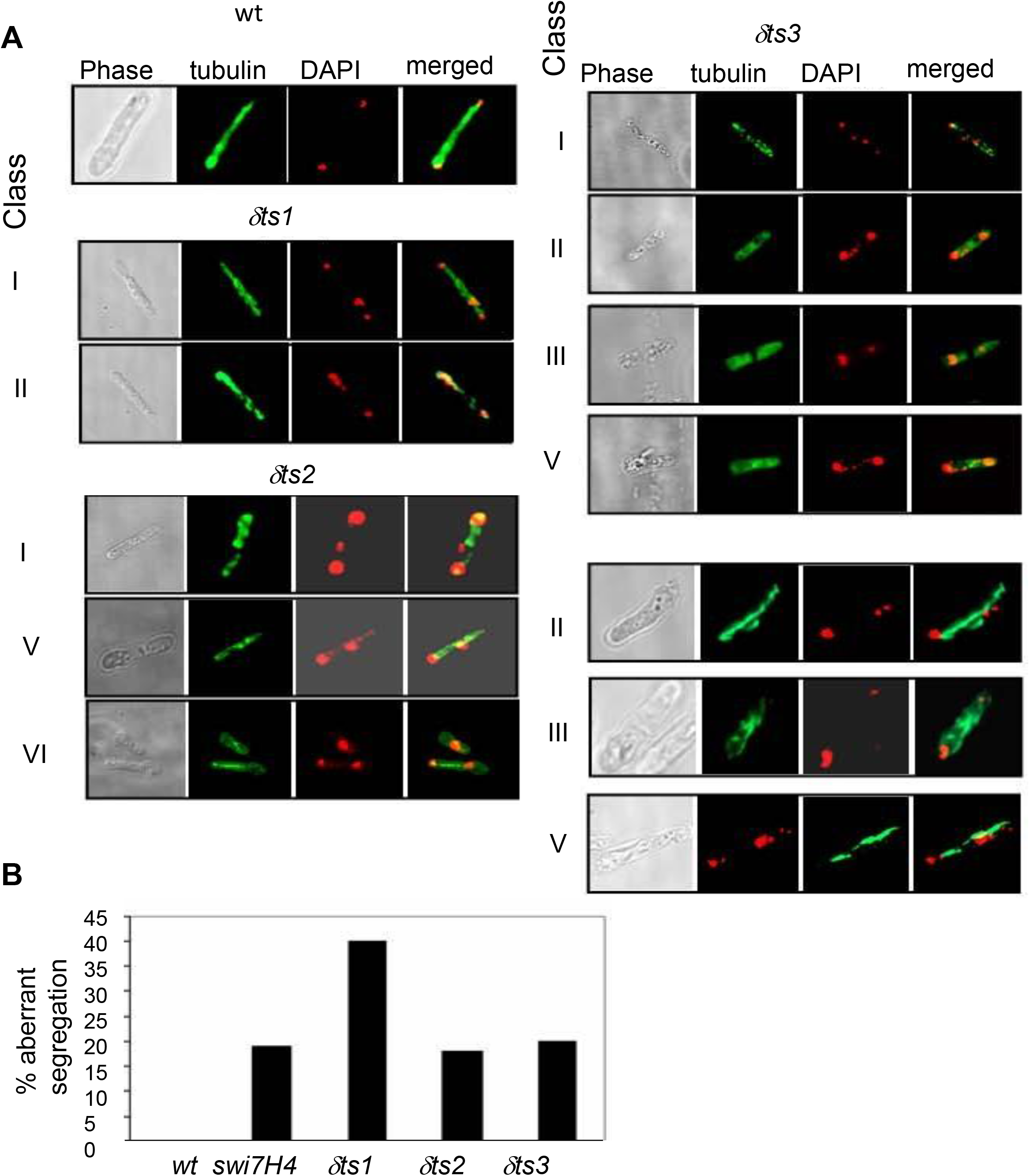
Chromosome segregation defects during mitosis in *polα* and *polδ* mutants. (A) Log phase cultures of wild type and *polα* and *polδ* mutant strains were subjected to DAPI staining and immunofluorescence microscopy using anti α-tubulin antibody. Representative images of the cells showing staining with anti α-tubulin antibody and DAPI and their overlaps are shown. Various classes of segregation defects shown by the mutants based on supplementary Fig. 2 are shown. (B) Histogram showing the quantitation of the aberrant segregation defects. In each case at least a total of 200 dividing cells were counted.

**Figure 6.**
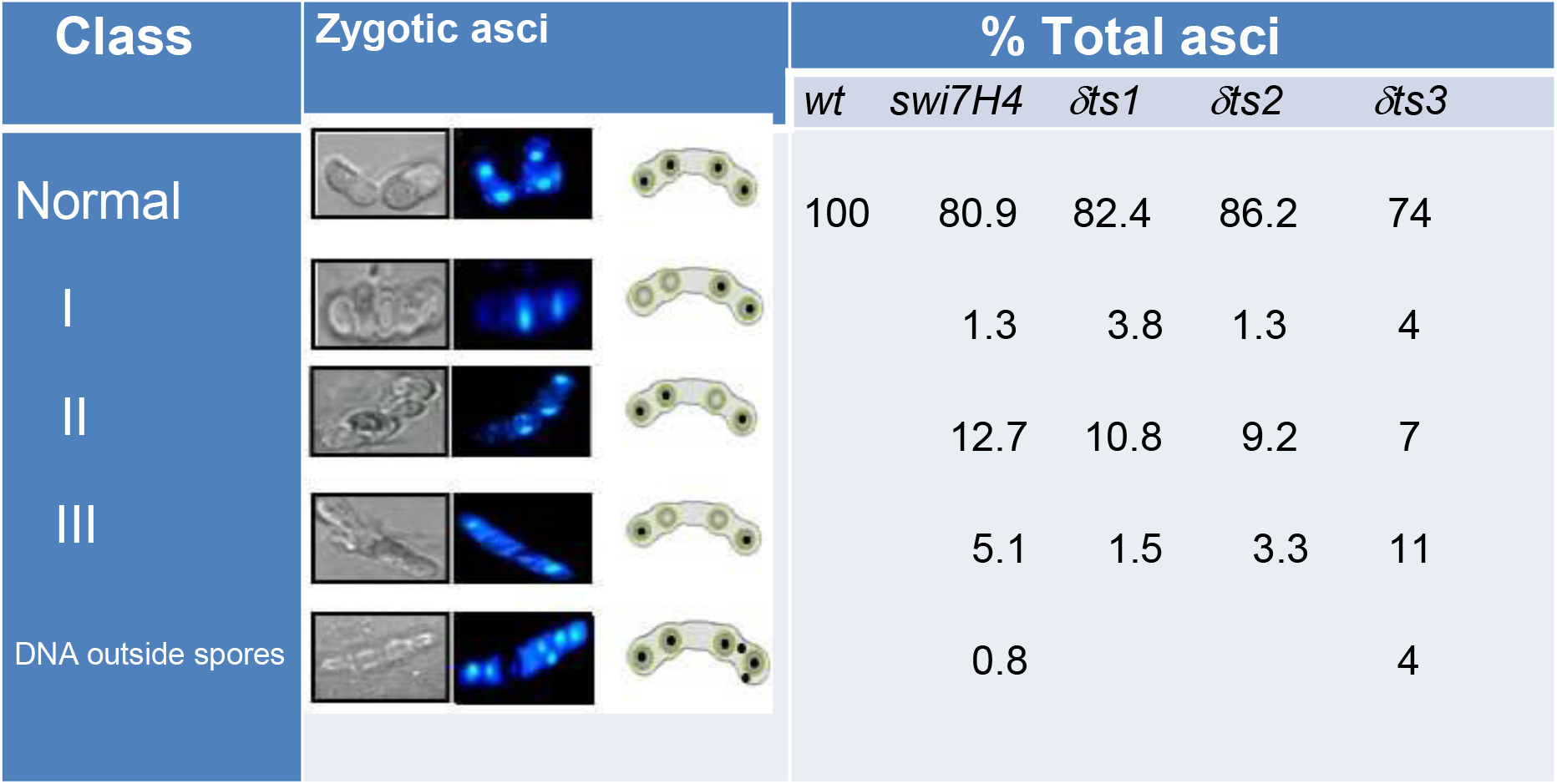
Meiotic segregation defects in *polα* and *polδ* mutants. Sporulating zygotic asci of wild type, *polα* and *polδ* mutant strains were stained with DAPI and visualized by confocal microscopy. Asci showing differential DAPI staining of zygotic ascospores were counted among a total of 200 asci and tabulated.

### Defects in meiotic chromosome segregation

Because proper chromatin compaction may be important for chromosome segregation during meiosis as well we checked whether Polα and Polδ, which have been shown above to be important for chromosome segregation during mitosis, are also required for proper chromosome segregation during meiosis. Therefore, we analyzed the phenotypes of 4-spored zygotic asci by monitoring their staining with DAPI to assess the segregation of DNA among the asci. Based on the different staining patterns, our results indicate that while the wild type asci show proper segregation of DAPI staining among all the four ascospores in all the asci, the mutants show reduced fidelity of segregation, with the defect being primarily at the second meiotic stage (Supplementary Figure 5).

## DISCUSSION

### A model for chromatin replication

In this study we have investigated a model for chromatin replication. We propose that DNA replication machinery may play an active role in replication of heterochromatin by virtue of direct interaction and recruitment of histone methyltransferase Clr4 and heterochromatin protein Swi6/HP1 on a replicating chromatin template. Our results show that indeed both DNA Polα and Polδ, the key enzymes involved in lagging and leading strand replication in all eukaryotes (25), bind to both Swi6/HP1 and Clr4 and are thus required for establishing the heterochromatin-specific histone modification code, i.e., methylation of histone H3 at Lys 9 position. This modification has been shown to be generally specific for heterochromatin regions, in contrast to enrichment of H3-Lys4 methylation in euchromatin regions, in not only in fission yeast but also higher eukaryotes (35,36). Histone H3 methylated at Lys9 is specifically bound by the chromodomain motif of Swi6/HP1 (37), a function again conserved during evolution. We speculate that the major components of DNA replication, namely DNA Polα and Polδ may recruit Swi6 and Clr4 on the replicating heterochromatin template and coordinate their cooperative actions-histone H3-Lys9 methylation, followed by binding by Swi6. Because of binding between Swi6 and Clr4, the process of methylation and binding by Swi6 may become highly cooperative and dynamic, leading to efficient duplication of the parental heterochromatin protein complement in the daughter chromatids. The coupling to the process of replication may have additional significance: the processive nature of replication may facilitate a rapid process of assembly of chromatin and ensure precision of duplication of the chromatin template in coordination with the S phase during cell cycle. Further, since the mechanism of replication, involving a coordinated function of DNA polymerases α and δ and other replication proteins and the proteins involved in heterochromatin assembly, is conserved during evolution, our model, suggesting a direct coupling of DNA replication machinery with active recruitment of heterochromatin proteins may also be conserved.

Consistent with this view, various findings in other organisms have highlighted the role of DNA replication components in silencing. For example, mutations in ORC suppress position-effect variegation (a hallmark of silenced regions wherein genes inserted near to silenced regions exhibit variegated expression patterns) in *Drosophila* (38). Mutations that disrupt silencing at rDNA and telomeres have been identified in DNA replication proteins and DNA-replication-related proteins such as DNA helicase Dna2p, PCNA loading factor Rfc1p (39,40). Also, mutations potentially suppressing silencing defects are found in DNA replication proteins including DNA Polε (41), the replication initiation factor Cdc45, PCNA and RF-C (42). Mutations in DNA Polα have also been shown to disrupt rDNA silencing and telomere-position effects in *S. cerevisiae* (43,44). The DNA replication-linked histone deposition factor, CAF1 (Chromatin Assembly Factor I) assembles histones specifically onto replicated DNA molecules via binding of the CAF1 p150 subunit to PCNA (44). Mutant forms of PCNA defective in CAF1 interaction are also defective in establishing silencing (45), suggesting that CAF1 and PCNA link DNA replication to chromatin assembly and silencing. Mutations in CAF1 and another chromatin assembly factor ASF1 also disrupt silencing (46,47). Mammalian CAF1 p150 subunit directly binds HP1 (48). PCNA associates with HDAC activity in human cells, strengthening the role of PCNA as a factor coordinating DNA replication and epigenetic inheritance (49). Recently, it has been shown that the methyl-CpG binding protein MBD1 couples histone H3 Lys9 methylation by SETDB1 to DNA replication and chromatin assembly (50).

Another important factor in coordinating DNA replication and heterochromatin assembly is the interaction between Swi6 and Clr4, as reported earlier (51), similar to interaction between mammalian Suv39H1 and mouse HP1 (52). Furthermore, genetic and molecular studies in *S. pombe* have shown a tightly coordinated spreading of Swi6 and the H3-Lys9-Me2 across the *K* region of mating type locus (53). In mammalian systems, co-localization and co-immunoprecipitation studies suggested an interaction between SUV39H1 and HP1 (54).A recent study analysed the interrelationships between histone H3-K9 methylation, transcriptional repression and HP1 recruitment and showed that targeting HP1 to chromatin required not only H3-K9 methylation but also a direct protein-protein interaction between Suv39H1 and HP1, underlining the importance of Swi6-Clr4 interaction in establishing heterochromatin states (55).

These studies, together with our results moot the question whether like replication, the process of heterochromatin replication is also semiconservative with the components of heterochromatin, like Swi6 and Clr4 segregating in a semi-conservative manner coincidentally with DNA replication. In a pertinent report Tagami *et al* showed that CAF1 forms a complex with a heterodimer of H3-H4 rather than the H3 -H4 tertramer, thus suggesting the possibility that H3-H4 dimer may be deposited at a replicating template, subsequent to segregation of two H3-H4 dimer halves along the segregating chromatids (56). Thus, the present results suggest that the Clr4-Swi6 complex may together be recruited by the DNA polymerases and help in carrying out a chromatin template-dependent copying of the H3-Lys9 methylation mark-coupled to Swi6 binding on the daughter chromatids in a highly processive manner. The nascent assembly of nucleosomes is executed by CAF1 (57). Since CAF1 also binds HP1 (58), Swi6 deposition may start with CAF1. Likewise, the binding of Swi6 to CAF1 (and Polα) may involve the chromoshadow domain of Swi6 and MOD1-interacting region (MIR; 59), which is present in CAF1 (59), Polα (17) and TIF1-α and TIF1-β (60; sequences similar to MIR are also found in Polδ). Once Swi6 is recruited, the Clr4 associated with it may bring about H3-Lys9 methylation and through positive feedback loop involving the interaction with Swi6, spread the region of heterochromatin in a catalytic and highly cooperative manner. Importantly, the interaction with DNA Polymerases would make the process of reassembly of heterochromatin highly cooperative and fast, which is important considering the narrow time window available for replication of heterochromatin region in *S. pombe* and metazonas, in general. In agreement, we find nearly 3-fold increase in the ratio of Swi6 to Polα during S phase, coinciding with the timing when heterochromatin region is replicated (61, 62; Balveer Singh, unpublished data). It would be interesting to study the role of DNA replication machinery vis-à-vis the chromatin assembly factor CAF1 in recruitment of Swi6 and Clr4.

However, these findings are difficult to reconcile with the results from budding yeast showing that silencing could be established on circular DNA molecules containing the mating type locus in the absence of DNA replication although, notably, passage through S phase was still essential (11,12). Since presence of the cis-acting silencer has been shown to be continuously required for heterochromatin in the budding yeast (63), some molecular event during S phase that recruits the heterochromatin protein Sir1 to the cis -acting silencer may substitute the process of replication. Alternatively, the mechanisms of silencing in the budding and fission yeast may be fundamentally different.

It remains to be seen whether, apart from Swi6 and Clr4, other silencing factors are also associated with DNA polymerases. Our preliminary results indicate a genetic interaction of *polα/swi7H4* mutant with all the silencing mutants *clr1-clr4* and *swi6*. Homothallic switching strains of these mutants as well as *swi7H4* mutant show nearly normal iodine staining as well as the level of switching, as measured by the percentage of zygotic asci, double mutants with *swi7H4* showed a drastic decrease in the level of sporulation as well as iodine staining (Supplementary Figure 6). These results suggest that together with Polα, the heterochromatin proteins may play a synergistic role in mating type switching as well.

### A role of DNA replication machinery in sister chromatid cohesion

It is also interesting to note that the heterochromatin protein assembly is coupled to replication on one hand and to the process of cohesion between sister chromatids on the other, which, in turn, ensures stable higher order organization, integrity of chromosomes and their faithful segregation during mitosis and meiosis. Our results suggest that DNA replication is also integrated to the latter processes possibly through its role in heterochromatin assembly. The process of DNA replication being processive, the heterochromatin assembly can be executed with high fidelity and speed, which in turn may recruit Cohesin in a highly cooperative manner. Accordingly, the process of chromatin assembly with cohesion being recruited both along the leading and lagging strand templates, can ensure a duplicative recruitment of Cohesin on both the daughter chromatids. This view is consistent with a recent report (64), which suggests recruitment of two separate cohesion complexes on the daughter chromatids. This speculative scenario is visualized in Figure 7. Consequently, in the replication mutants even a slightly reduced binding to the heterochromatin components may elicit severe consequences including defects not only in silencing, which is a functional correlate of the heterochromatin, but also in the reproduction of higher order chromosome structure and recruitment of Cohesin, leading to enhanced chromosome loss, chromatid segregation defects, aneuploidy and loss of viability. Thus, our studies indicate a coordination and integration of the process of DNA replication with heterochromatin assembly as well as sister chromatid cohesion. Because of the conservation of these machineries during evolution, eukaryotic species may have a highly conserved system of spatio-temporal coordination of the replication machinery with heterochromatin assembly and sister chromatid cohesion. Any defects in these steps may lead to aberrant chromosome segregation eliciting disease states including cancer.

**Figure 7.**
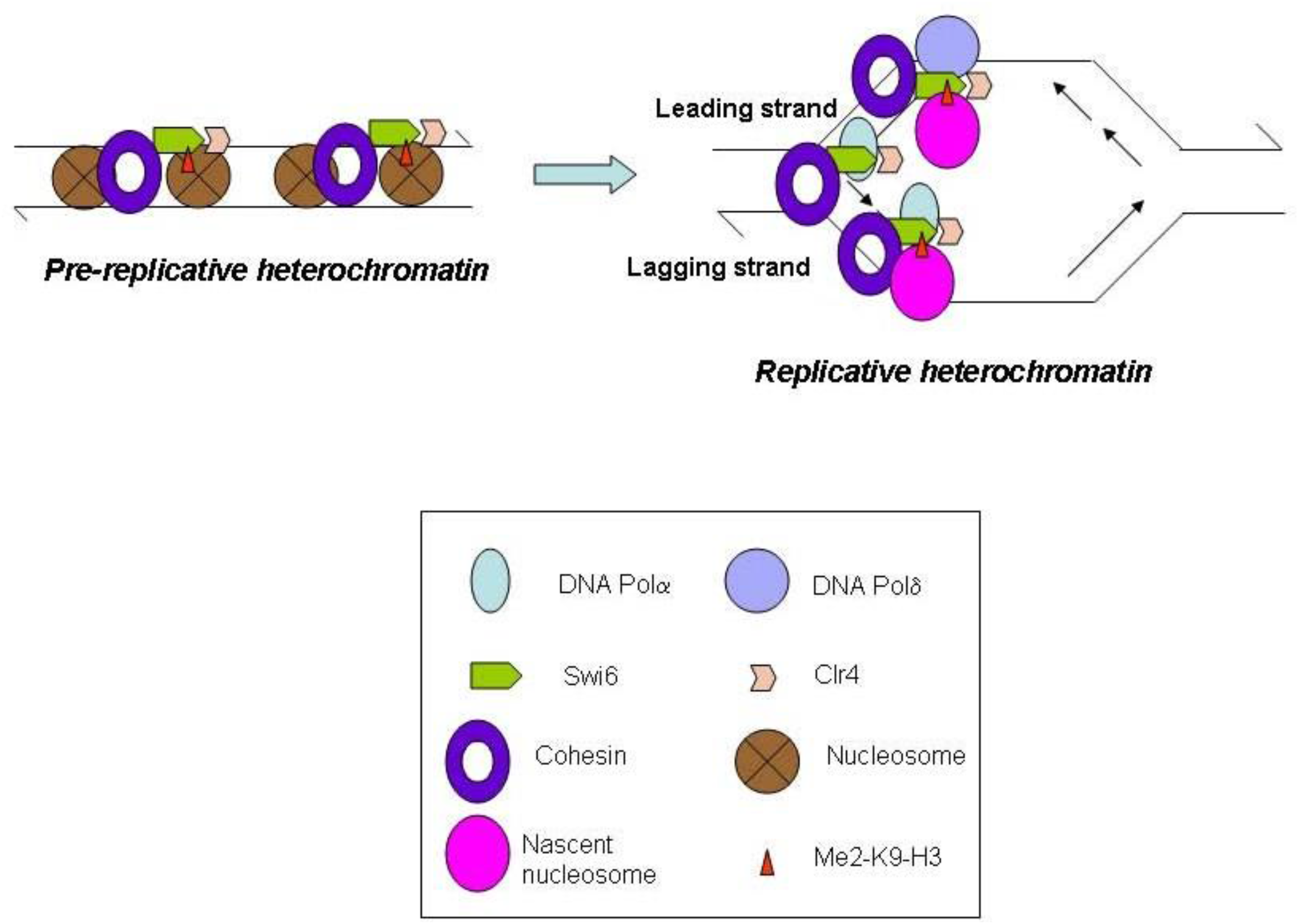
Chromatin replication model. A speculative model depicting the coupling of DNA replication machinery comprising of DNA Polα and Polδ to heterochromatin proteins Swi6/HP1 and Clr4, and through Swi6, with cohesion, thus bringing about a replication assisted duplication of the epigenetic information of heterochromatin and sister chromatid cohesion.

## Acknowledgements

We are grateful to Teresa Wang for the *polδ* mutants, H. Okayama for the *swi7H4* mutant, G. Thon for the strain PG1649, R. Allshire for the strains carrying *ura4* and *ade6* reporters at centromere, K. Ekwall for the strains HU393 and HU395, A. Pidoux for the plasmid harbouring GFP-Swi6, J.P. Javerzat for the strain harbouring HA-tagged rad21. M. Yanagida fro strain having HA- and GFP-tagged cnp1 and S. MacNeil for the plasmid expressing GST-tagged Polδ. We thank S. Haldar and A. Sharma for reading the manuscript. This work was supported by the Intramural support from Council of Scientific and Industrial Research (CSIR), New Delhi, India. Kamlesh Bisht and Nandni Nakwal are recipients of Senior Research Fellowship from CSIR.

