## Supplementary file for "Heterochromatin Replication: Direct Interaction of DNA replication machinery with heterochromatin code writer Clr4/Suv39 and reader Swi6/HP1 in *S. pombe*"

#### Supplementary Section

##### Supplementary methods

**Immunofluorescence method.** Immunofluorescence with TAT1 antibody staining was done as follows. Cells were fixed with methanol at -20°C for 10 min to overnight. Fixed cells were pelleted by centrifugation (6000 rpm for 2 min), resuspended in 1ml of PEM buffer and transferred to an eppendorf tube, followed by three washings with PEM (1ml per wash). Next, the cells were washed once in PEMS buffer (PEM + 1.2M Sorbitol) and cell walls were digested by 1mg/ml zymolyase 100T (dissolved in PEMS) for 15-30 minutes at 37°C. Incubation was carried out until cell walls of about 60% of the cells were digested. Cell wall digestion was monitored under a microscope by adding 1μl SDS to 9μl cell suspension. Following digestion, cells were washed thrice with PEMS and permeabilised by incubation in 1ml PEMS containing 1% Triton X-100 for 30 sec. Cells were next washed three times in PEM buffer and resuspended in 1ml PEMBAL buffer (PEM buffer with 1% BSA, 0.1% sodium azide and 100mM lysine hydrochloride). Cells in PEMBAL were incubated for at least 30 min at RT with gentle rocking. Cells were recovered by centrifugation, resuspended in 30μl of diluted mouse monoclonal α-tubulin antibody (Sigma) in PEMBAL (2μl antibody in 28μl PEMBAL). Antibody incubation was done for 16hrs at RT with rotation followed by three washings with PEMBAL buffer. The stained cells were resuspended in 100μl of diluted anti-mouse FITC conjugated antibody (Santacruz) in PEMBAL (1μl antibody + 99μl PEMBAL) and incubated for 4-16 hours at RT with rotation. Tubes were covered with aluminum foil during incubation as FITC is light-sensitive. Cells were again washed thrice with PEMBAL and finally resuspended in 100μl PEMBAL. 4-5μl of the stained cells were mixed with 4μl of 4X DAPI and analysed using Carl-Zeiss Fluorescence microscope.

#### Supplementary Figure Legends

**Supplementary Figure 1.** ChIP-on-ChIP assay to quantitate the level of H3-Lys9-Me2 and Swi6 in wild type versus *swi7H4*/*polα* and *polδts2* mutants at the *cenII*. Double vertical arrows indicate the regions in the outer repeat reduction in the level of H3-Lys9-Me2 and Swi6.

**Supplementary Figure 2.** ChIP-on-ChIP assay to quantitate the level of H3-Lys9-Me2 and Swi6 in wild type versus *swi7H4*/*polα* and *polδts2* mutants at the *cenII*.

**Supplementary Figure 3.** Delocalization of Swi6 oin *polδ* ts mutants. (A) Confocal microscopy results quantitating the number of cells having 1, 2, 3 or >3 spots of GFP-Swi6 in wt and *polδts* mutants. (B) *polα* and *polδ* mutants display enhanced sensitivity to cold and the microtubule destabilizing drug thiabendazole. Equivalently grown cultures of wild type, *polα* and *polδ* mutant strains were taken and 5μl of 10 -fold serial diutions of the cultures were spotted on normal complete plates and plates containing thiabendazole (TBZ) . While the normal plates were incubated at 25$^{O}$C and 18$^{O}$C, the plate containing TBZ was incubated at 25$^{O}$C for 4-5 days.

**Supplementary Figure 4.** Reduced sister chromatid cohesion in *swi7H4* and *δts2* mutant. (A) Cells of log phase cultures of wild type, *polα* and *polδ* mutant strains harbouring LacI-GFP fusion protein, and the LacO DNA repeat inserted at the *lys1* locus linked to centromere 1 (*cen*1-GFP), were visualized by confocal microscopy. A total of 200 cells were counted and those containing one and two spots of GFP were counted and shown in the histogram. (B) Reduced localization of Cohesin subunit Rad21 at *cenI* in *polα* and *polδ* mutants. Wild type, *polα* and *polδ* mutant strains harbouring HA-tagged *rad21* gene were subjected to ChIP assay and the relative localization of Rad21 at *cenI* (*dh*) and *act1* was quantitated. Numbers denote the *dh*/*act1* ratio of the immunoprecpitated (IP) versus whole cell extract (WCE) samples.

**Supplementary Figure 5.** Schematic diagram depicting the normal and mutant pattern of segregation of nuclei during the first and second meiotic stage.

**Supplementary Figure 6.** Cumulative effect of *swi7H4* with heterochromatin protein mutants *swi6*, *clr1-clr4* on efficiency of mating type switching. Single and double mutants of *swi7H4* and *swi6*, *clr1-clr4* were streaked on pombe minimal plates. After 4-5 days of growth at 25ºC, the colonies were stained with iodine and photographed. In parallel, cells of the colonies were examined uner light microscope and the level of switching quantitated by using the equation. % switching: [2(no. of zyngotic asci)/haploid cells + 2 (no. of zyngotic asci)] x 100.

**Table 1.** List of strains used in this study

| Strain Name | Genotype | Source |
|---|---|---|
| FY2002 | $h^+$ leu1-32 ura4DS/E ade6DN/N imr1L-Nco::ura4 otr1R-Sph::ade6 | R. Allshire |
| SPJ834 | $h^+$ leu1-32 ura4D18 ade6-216 imr1L-Nco::ura4 otr1R-Sph::ade6 δts1 | This study |
| SPJ838 | $h^+$ leu1-32 ura4DS/E ade6DN/N imr1L-Nco::ura4 otr1R-Sph::ade6 δts2 | This study |
| SPJ839 | $h^+$ leu1-32 ura4DS/E ade6DN/N imr1L-Nco::ura4 otr1R-Sph::ade6 δts3 | This study |
| FY336 | $h^+$ leu1-32 ura4DS/E ade6DN/N cnt1/TM1-Nco::ura4 | R. Allshire |
| SPC20 | $h^+$ leu1-32 ura4DS/E or ura4D18 ade6DN/N cnt1/TM1-Nco::ura4 δts3 | This study |
| SPC28 | $h^+$ leu1-32 ura4DS/E or ura4D18 ade6DN/N cnt1/TM1-Nco::ura4 δts1 | This study |
| SPC29 | $h^+$ leu1-32 ura4DS/E or ura4D18 ade6DN/N cnt1/TM1-Nco::ura4 δts2 | This study |
| PG1649 | mat1P 17::LEU2 REIII mat3M::ade6 oriI leu1-32 ura4D18 ade6-210 | G. Thon |
| FY501 | $h^+$ leu1-32 ura4DS/E ade6DN/N imr1L-Nco::ura4 | R. Allshire |
| SPC14 | $h^+$ leu1-32 ura4DS/E or ura4D18 ade6DN/N imr1L-Nco::ura4 δts1 | This study |
| SPC15 | $h^+$ leu1-32 ura4DS/E or ura4D18 ade6DN/N imr1L-Nco::ura4 δts2 | This study |
| SPC16 | $h^+$ leu1-32 ura4DS/E or ura4D18 ade6DN/N imr1L-Nco::ura4 δts3 | This study |
| SP1137 | $h^{90}$ mat2::ura4 ura4D18 ade6-210 | A. Klar |
| SPJ541 | $h^{90}$ leu1-32 his2 ura4 D18 ade6–216 | This study |
| SPC11 | $h^{90}$ leu1-32 his2 ura4 D18 ade6–216 δts1 | This study |
| SPC18 | $h^{90}$ leu1-32 his2 ura4 D18 ade6–216 δts2 | This study |
| SPC19 | $h^{90}$ leu1-32 his2 ura4 D18 ade6–216 δts3 | This study |
| KSP3 | Msmto ura4DS/E ade6DN/N imr1L::ura4.otrR:sph::ade6 δ ts2 | This study |
| SPA300 | Msmto leu1-32 ura4DS/E otr1R(Sph1):: ura4 swi7H4 his2 ade6DN/N | This study |
| FY521 | $h^+$ Ch16 m23::$ura4^+$ ura4DS/E ade6-210 (ch16 ade6-216) | R. Allshire |
| SPC44 | $h^+$ Ch16 m23::$ura4^+$ ura4DS/E ade6-210 (ch16 ade6-216), δts1 | This study |
| SPC42 | $h^+$ Ch16 m23::$ura4^+$ ura4DS/E ade6-210 (ch16 ade6-216), δts2 | This study |
| SPC43 | $h^+$ Ch16 m23::$ura4^+$ ura4DS/E ade6-210 (ch16 ade6-216), δts3 | This study |
| HU393 | $h^+$ leu1/YIP 2.4 pUC $ura4^+$-7 ura4DS/E leu1-32 ade6M216 | K. Ekwall |
| SPJ990 | $h^{90}$ leu1-32 ura4 D18 swi6 ::ura4 ade6-216 | This study |
| SPC39 | $h^{90}$ leu1-32 ura4 D18 swi6 ::ura4 ade6-216 δts1 | This study |
| SPC40 | $h^{90}$ leu1-32 ura4 D18 swi6 ::ura4 ade6-216 δts2 | This study |
| SPC41 | $h^{90}$ leu1-32 ura4 D18 swi6 ::ura4 ade6-216 δts3 | This study |
| SPJ695 | $h^{90}$ leu1-32 ura4 D18 mat3M::ura4 ade6-216 | This study |
| SPS180 | $h^{90}$ leu1-32 his2 ura4 D18 ade6–210 swi6 ::ura4 nmt1GFP-swi6 | This study |
| SPS184 | $h^{90}$ leu1-32 ura4 D18 cut4-553 ade6-210/ pREP3X | This study |
| SPS185 | $h^{90}$ leu1-32 ura4 D18 cut4-553 ade6-210 intHA-cut4 | This study |
| SPA236 | Msmto leu1-32 ura4 D18 REII mat2::ura4 ade6-216 | Ahmed et al 2001 |
| SPJ633 | Msmto leu1-32 ura4 D18 REII mat2::ura4 ade6-216 δts2 | This study |
| SPC22 | Msmto leu1-32 ura4 D18 REII mat2::ura4 ade6-216 δts1 | This study |
| SPC24 | Msmto leu1-32 ura4 D18 REII mat2::ura4 ade6-216 δts3 | This study |
| FY336 | $h^+$ leu1-32 ura4 DS/E cnt1/TM1 {NcoI}:: ura4 ade6-210 | R. Allshire |
| FY501 | $h^+$ leu1-32 ura4 DS/E imr1L-Nco::ura4 ade6210 | R. Allshire |
| 2383 | $h^+$ lacO-lys1+GFP-lacI-NLS-$his7^+$ | M. Yanagida |
| SPC57 | $h^+$ leu1-32 ura4D18 lacO-lys1+GFP-lacI-NLS-$his7^+$ δts1 | This study |
| SPC58 | $h^+$ leu1-32 ura4D18 lacO-lys1+GFP-lacI-NLS-$his7^+$ δts2 | This study |

### Supplementary Figure 1

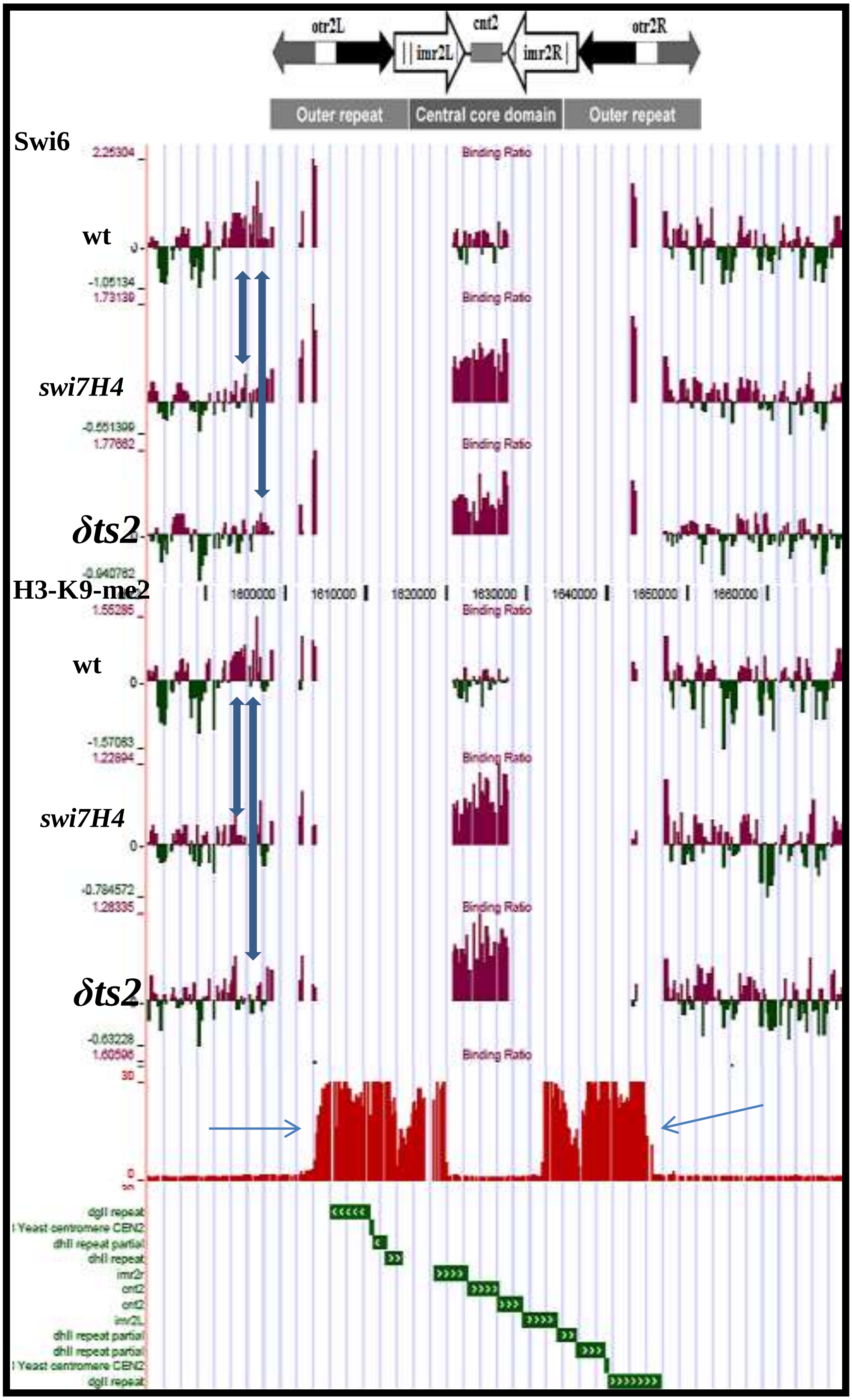

Supplementary Figure 2

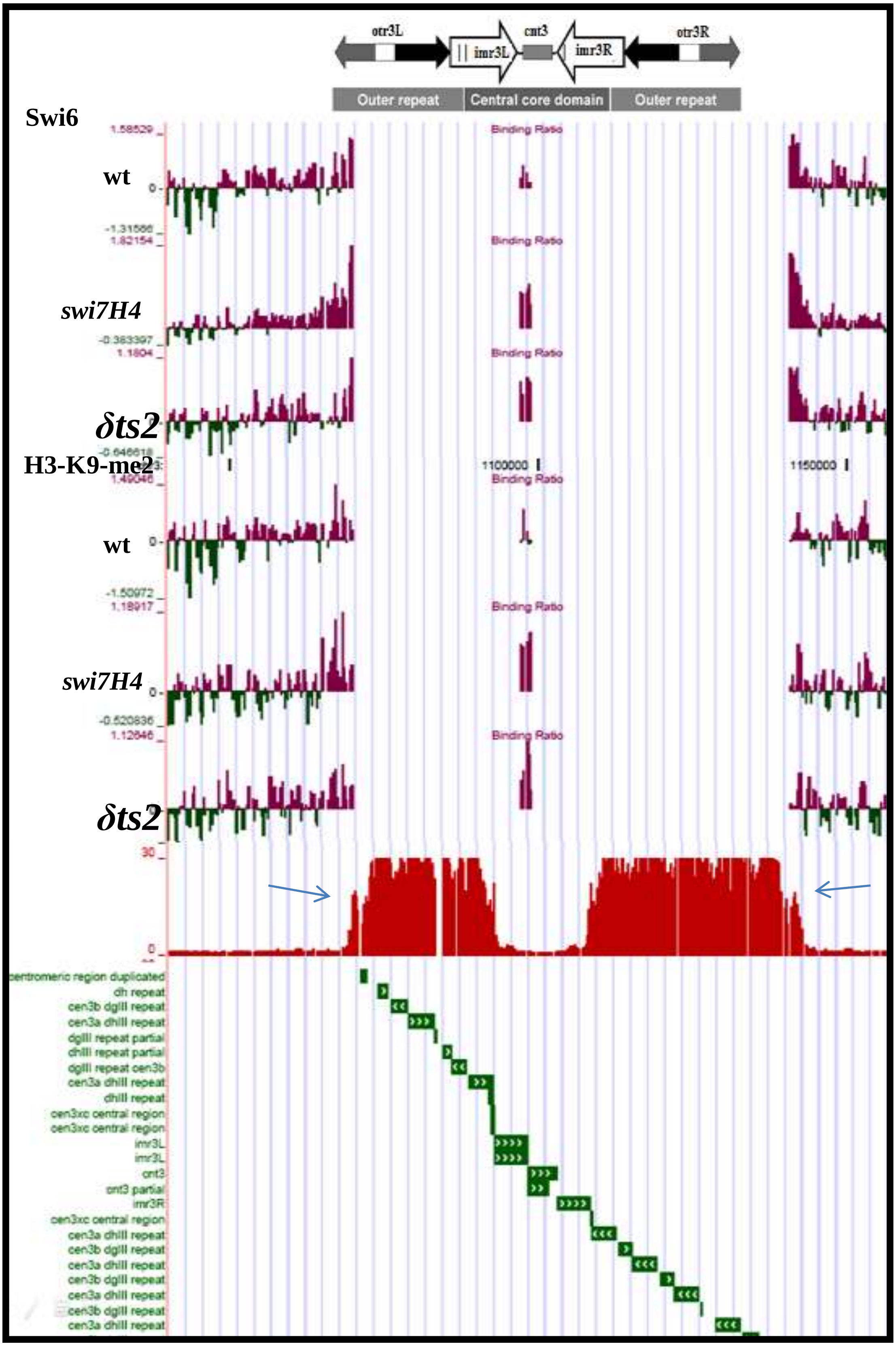

Supplementary Figure 3

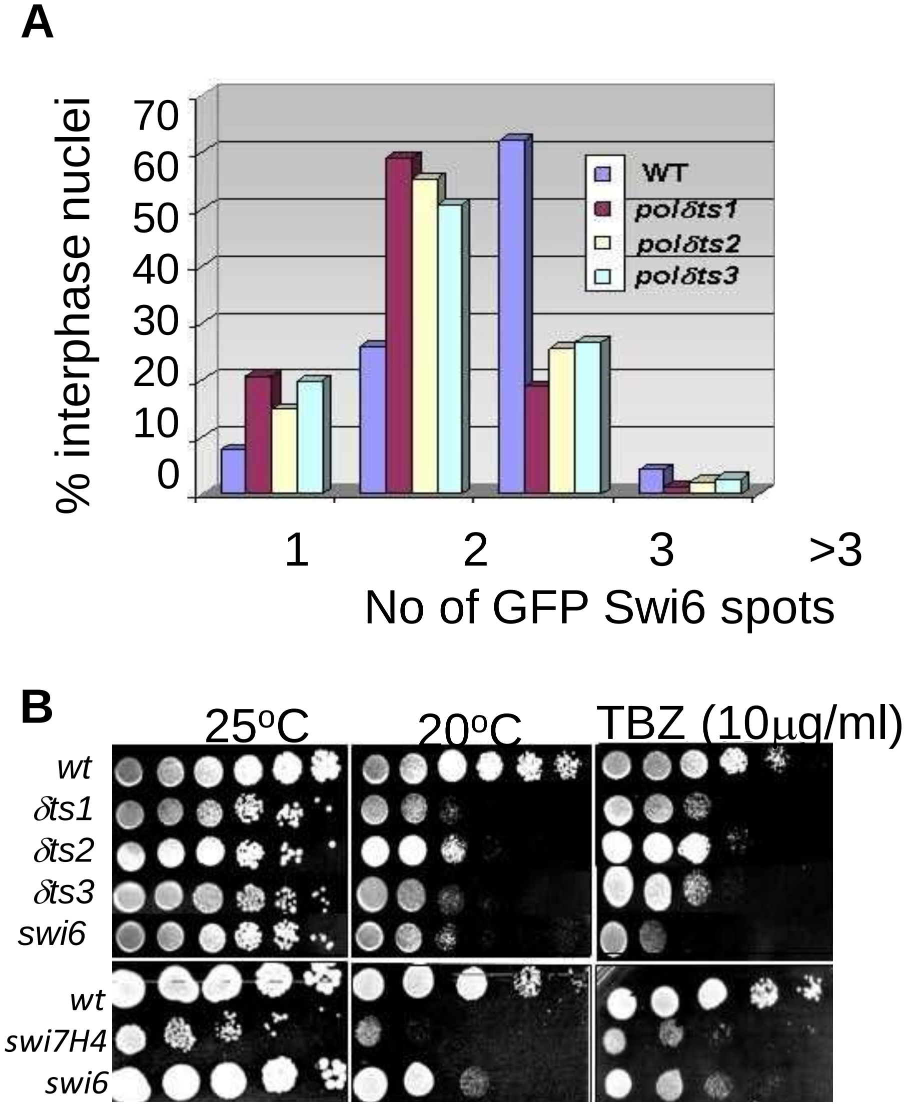

Supplementary Figure 4

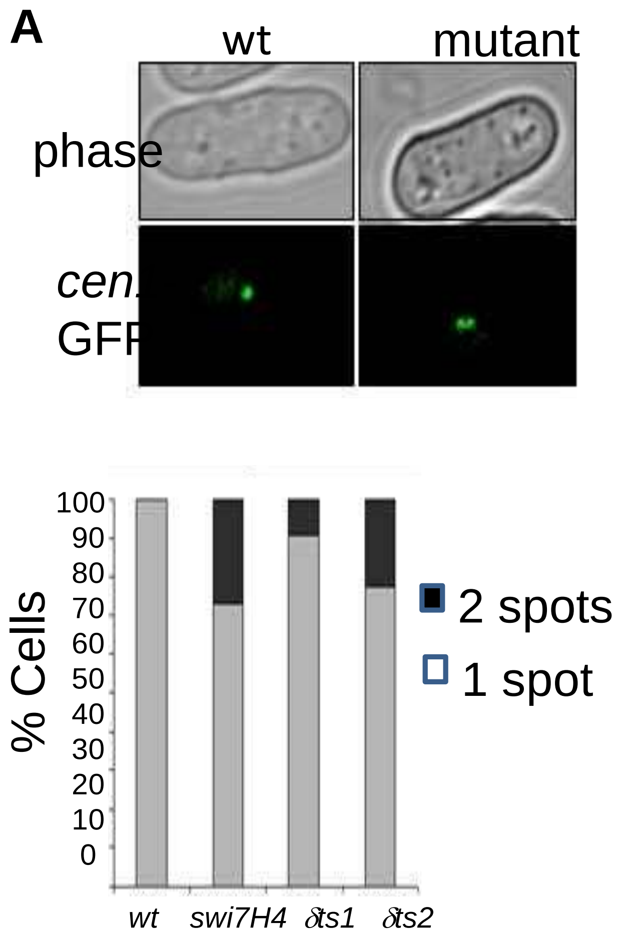


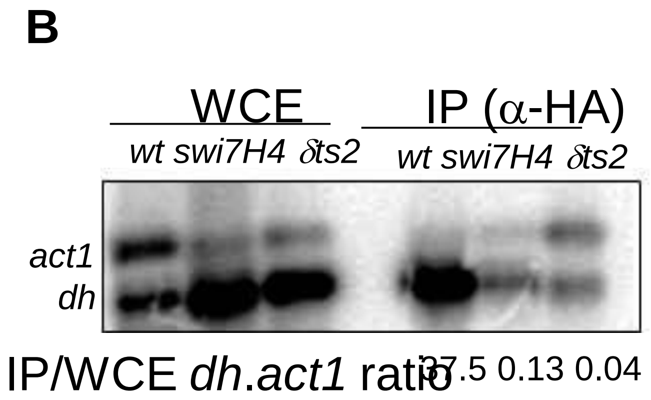

### Supplementary Figure 5

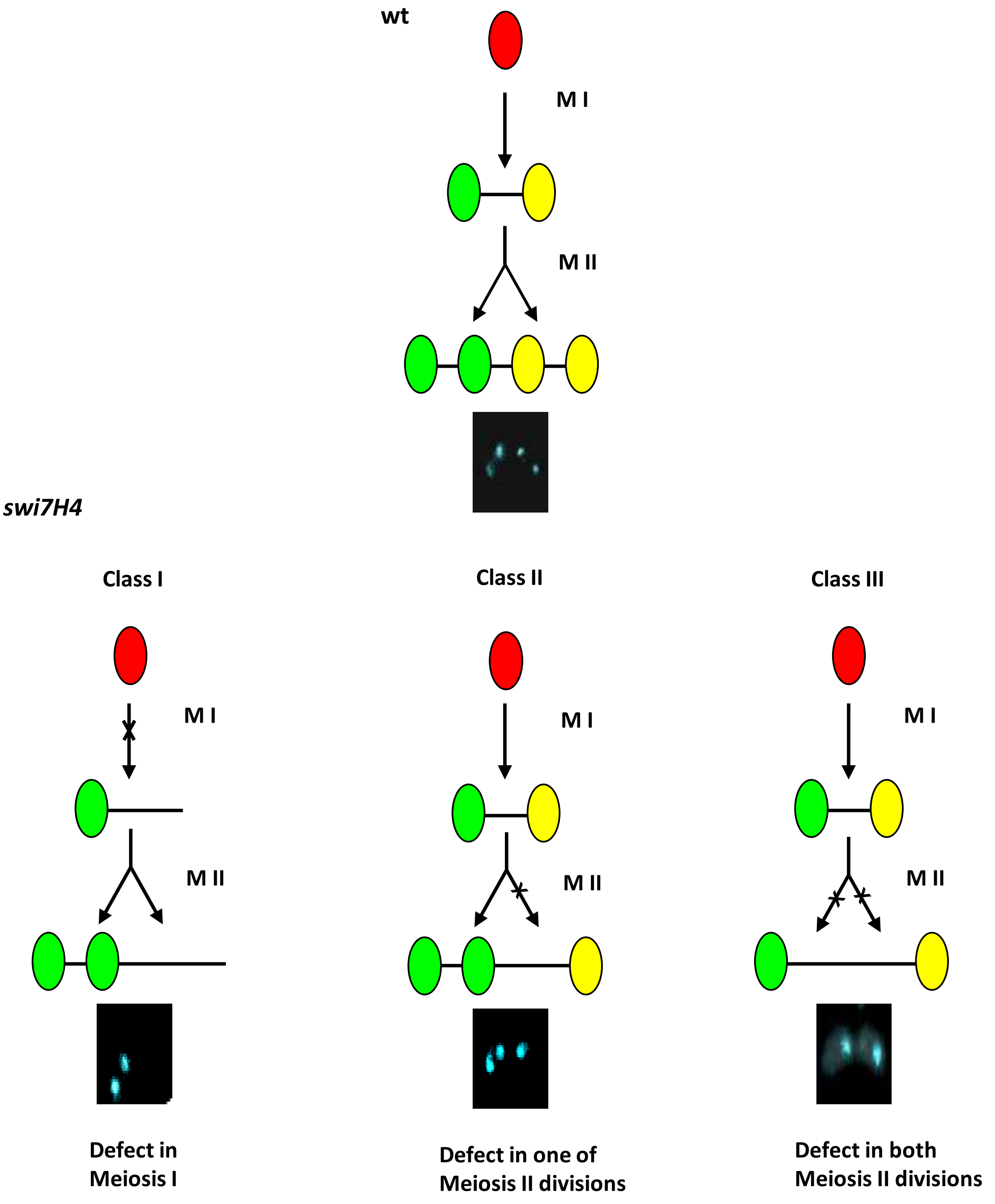


**Fig. 3.3.4B Diagrammatic representation of the classes of meiotic defects observed in *swi7H4* mutant**

Upper panel shows the normal meiotic division in *wt* strain. Lower panels depict the meiotic defects observed in *swi7H4* mutant. Below each sketch, a representative sporulation pattern of each class is shown.

Supplementary Figure 6

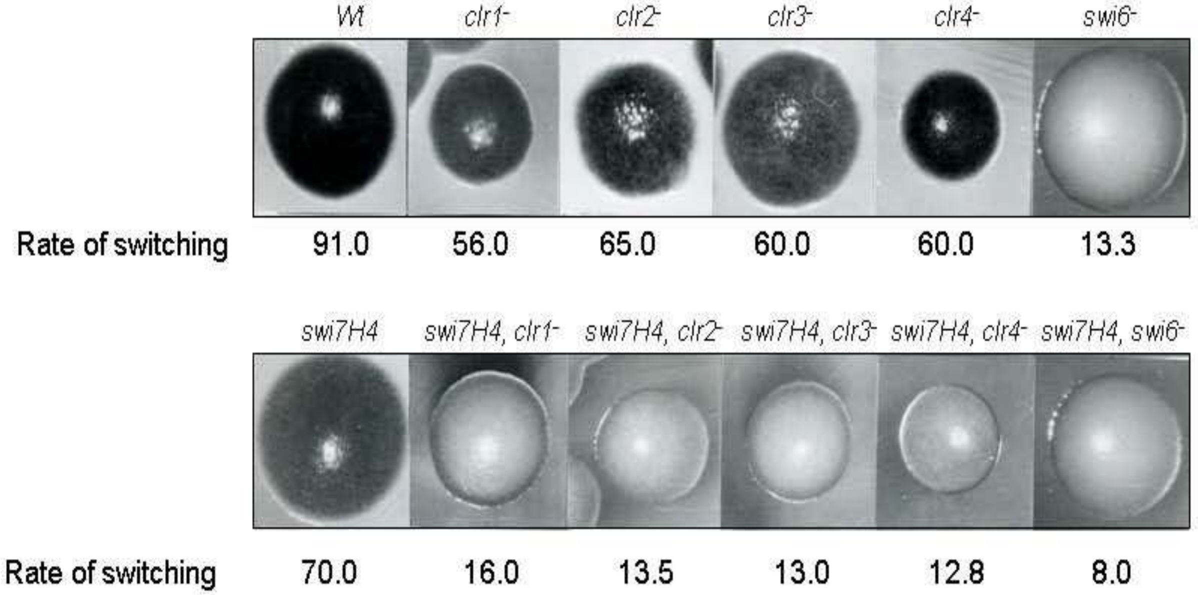
